# Impacts of projected elevated temperature on estimated chilling accumulation and bud break of rabbiteye blueberries

**DOI:** 10.1101/2025.07.08.663765

**Authors:** Lovely Mae F. Lawas, Giovani Rossi, Melba R. Salazar-Gutierrez, Di Tian, Courtney P. Leisner

## Abstract

Climate change poses a significant threat to perennial fruit crops, including blueberries, by altering temperature-dependent developmental processes such as dormancy and bud break. This study investigates the projected impacts of mid-21st century warming on chilling accumulation and phenological development in rabbiteye blueberry (*Vaccinium virgatum*) cultivars ‘Krewer’ and ‘Titan’. Using downscaled climate projections from 20 global climate models under the RCP 8.5 scenario, we estimated future temperature trends for a key southeastern U.S. blueberry production region. We quantified changes in chilling hours, chilling portions, and freeze event probabilities between historical (1981–2000) and projected (2041–2070) periods. Results indicate a substantial reduction in chilling accumulation—chilling hours decreased by 66.6% and chilling portions by nearly half—alongside a delayed onset and shortened duration of chilling periods. Despite warming, freezing events are still expected during critical phenological windows, potentially increasing frost damage risk. Growth chamber experiments simulating ambient and elevated temperature regimes revealed significant cultivar-specific shifts in the timing and rate of bud break and bloom stages. Logistic models developed from these data provide predictive tools for assessing phenological responses under future climates. These findings highlight the vulnerability of rabbiteye blueberry to climate-induced dormancy disruption and underscore the need for adaptive management strategies and cultivar selection to sustain productivity in a warming climate.

## 1. Introduction

The observed global warming over the past century is predicted to continue in the future, with an estimated increase in global surface temperature of 1.5 °C in the near term (2030-2035), and by 1.4 – 4.4 °C towards the end of the century (IPCC, 2023). This future climate change poses a threat to crop production (IPCC, 2023) as temperature is a key factor affecting plant growth and development (Hatfield & Prueger, 2015). Plant responses to extreme and moderately high temperatures have been well documented in annual crops such as cereals (e.g., barley, maize, rice, wheat) and legumes (e.g., bean, pea, soybean) (Zhu et al., 2021; Sita et al., 2017). In comparison, there is limited information known about the impacts of elevated temperatures on perennial fruit crops (Leisner, 2020).

Understanding how climate change impacts perennial plants is of importance as predicted changes in seasonal temperatures cause unique vulnerabilities in these plants (Winkler et al., 2016). For example, springtime freeze risk is a key climate-influenced vulnerability of perennial plants (Winkler et al., 2016). The growth of perennial plants is also impacted by elevated temperatures, as these plants live more than one year, and most cycle between growth and reproduction over multiple seasons (Friedman, 2020). To withstand unfavorable growth conditions during seasonal environmental changes, perennial plants located in temperate climates enter a dormancy phase which requires specific temperature and photoperiod (day length) cues to resume growth in the spring (Luedeling, 2012). Improper timing of flowering due to ineffective winter dormancy and the subsequent impacts on fruit production have significant additional consequences on global food security (Fadón & Rodrigo, 2018; Luedeling, 2012). Therefore, is it important to understand how global warming impacts dormancy traits in agriculturally important perennial plant species.

Blueberry (*Vaccinium spp.*) is a perennial shrub native to North America and grown in many parts of the world (Hancock et al., 2008). Its fruits are known for their presence of bioactive compounds including vitamins, minerals, and phenolic compounds such as flavonoids, phenolic acids, and anthocyanins (Nile & Park, 2014; Rossi et al., 2022). These compounds have positive effects on human health, including acting as antioxidants, preventing diseases, and containing antimicrobial properties (Shen et al., 2014). Aside from their health benefits, blueberries are an economically important fruit crop, with a global gross production value of $2.85 million in 2022 (FAO, 2024).

One important feature of blueberry cultivation is the presence of ecotypes determined by their temperature requirements for growth and development. In the U.S., most of the blueberry production involves the northern highbush ecotype (*V. corymbosum*), which are adapted to cooler temperatures. In warmer regions like the southeast U.S., southern highbush (*V. corymbosum* with introgression of *V. darrowii*) and rabbiteye (*V. virgatum* Ait. [syn. *V. ashei* Reade]) ecotypes are predominantly grown (Hancock et al., 2008). Most rabbiteye cultivars are pure *V. ashei*, which is known for heat tolerance (Edger et al., 2022), but rabbiteye ecotypes still show vulnerability high temperatures (Yang et al., 2019).

There are many ways in which blueberry development and productivity can be negatively impacted by heat stress (Chen et al., 2012; Zheng et al., 2017; Hao et al., 2019). The predicted increase and fluctuations in temperature under future climates may impact plant growth and photosynthetic activity (Salazar-Gutiérrez et al., 2023), and elevated temperatures can also impair proper development of flowers and fruits (Lobos & Hancock, 2015) by limiting the plant’s ability to acquire sufficient chilling hours which are needed for optimum flower and fruit development (Cifuentes-Carvajal et al., 2023; Drogudi et al; 2023). Subsequent early bud break and flowering may lead to yield losses due to spring frost injury (Gao et al., 2019). Additional work is needed, therefore, to understand how elevated temperature impacts chilling accumulation, dormancy, and budbreak in blueberry.

This study aimed to empirically determine the impacts of projected elevated temperature on chilling accumulation and key dormancy phenotypes in blueberry plants grown under realistic future climate conditions. To do this we analyzed changes in chilling hours and freeze events for blueberry cultivars predicted for the southeast U.S. growing region using downscaled global climate model outputs for the mid-21^st^ century. Next, we modeled the impact of temperature predicted for the mid-21^st^ century on bud break and bloom in commercially grown fresh market rabbiteye cultivars using controlled growth chamber experiments. We then analyzed the temperature effect on completion of the bud break stage in blueberries for future modeling.

Findings from this work will help identify key impacts of temperature on budbreak in blueberry and quantify the changes in chilling hours and freeze events for blueberry cultivars predicted for the mid-century in the southeast U.S.

## 2. Materials and Methods

### 2.1. Sources of climate data for projected future temperature determination

To accurately determine the impacts of elevated temperature on key floral stages of blueberry after budbreak (early bloom and late bloom) downscaled global climate model (GCM) data for the mid-century period (2041-2070) was used. GCM data was obtained from the Multivariate Adaptive Constructed Analogs (MACA) dataset (https://climate.northwestknowledge.net/MACA/index.php). The MACA method (Abatzoglou & Brown, 2012) is a statistical downscaling approach for bias correction of GCM outputs. It was used to downscale the model output of 20 GCMs from the Coupled Model Inter-Comparison Project 5 (Taylor et al., 2012). MACA data sets were obtained from the MACAv2-METDATA dataset https://climate.northwestknowledge.net/MACA/data_csv.php). To further downscale the GCM data for the southeastern U.S. the delta change factor method (Anandhi et al., 2011) was used. This method utilizes local weather station data to estimate future climate scenarios. Local weather data from Bacon County (Alma, GA, USA) was selected for downscaling. This location was chosen because it is a representative area in the southeastern U.S. for blueberry production, corresponding to 49% of the total production area in Georgia and 28% in the southeast (USDA NASS, 2022).

In this study, we used three data sets: Reference, MACA Reference, and MACA Projected. The Reference data set consisted of daily near-surface minimum (T_min_) and maximum (T_max_) temperatures from 01 January 1981 to 31 December 2000 downloaded from the weather station (ID # GHCND:USW00013870) located at the Alma Bacon CO Airport (NOAA National Centers for Environmental Information, 2022). MACA Reference consisted of monthly near-surface historical T_min_ and T_max_ temperatures from January 1981 to December 2000, while MACA Projected included data from the Representative Concentration Pathway (RCP) 8.5 scenario for the period 2041-2070. MACA Reference and MACA Projected data sets comprise data from all 20 GCMs. For each GCM, the gridded 1/24-deg MACAv2-METDATA data sets for T_min_ and T_max_ for the location at latitude 31.5084 and longitude −82.4513 were obtained.

### 2.2. Determining future projected temperatures for blueberry growing regions using downscaled climate models

To determine future projected temperatures for our selected blueberry growing region the raw data obtained from MACA and the weather station were processed by getting the average T_min_ and T_max_ on a daily (Reference) or monthly (MACA Reference and MACA Projected) basis across the years included (2041-2070 for MACA projected data and 1981-2000 for MACA Reference data and Reference data). The MACA Reference and MACA Projected data sets contain the monthly mean T_min_ and T_max_ for each of the 20 GCMs. The projected temperature change for the mid-century period (2041-2070) compared to the reference period (1981-2000) was then calculated as the delta (Δ) value.

The ΔT_min_ and ΔT_max_ for the projected period 2041-2070 was calculated using the following equations:

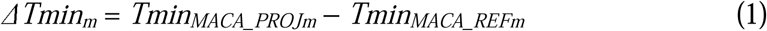

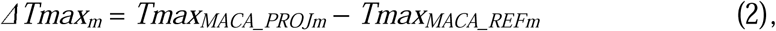

where Δ*Tmin_m_*and Δ*Tmax_m_* are the deltas of the average monthly minimum and maximum temperatures, respectively, for month *m*; *Tmin_MACA_PROJm_*and *Tmax_MACA_PROJm_*: average daily minimum and maximum temperatures, respectively, for month *m* (derived from the MACA dataset for RCP 8.5 scenario during the projected period); *Tmin_MACA_REFm_* and *Tmax_MACA_REFm_*: average daily minimum and maximum temperatures, respectively, for month *m* (derived from the MACA dataset for the historical period).

The deltas were then used to calculate the daily T_min_ and T_max_ based on the formula:

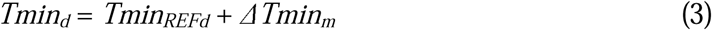

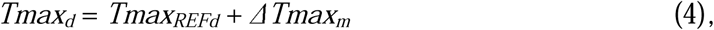

where *Tmin_d_* and *Tmax_d_*: minimum and maximum temperatures, respectively, for day *d* in month *m*; *Tmin_REFd_* and *Tmax_REFd_*: average minimum and maximum temperatures, respectively, for day *d* in month *m* (data from the reference period obtained from the weather station); Δ*Tmin_m_* and Δ*Tmax_m_*: deltas of the averaged daily minimum and maximum temperatures, respectively, for month *m*. Lastly, *Tmin_d_* and *Tmax_d_* were used to estimate hourly temperatures for the duration 01 January to 31 December for both the reference (1981-2000) and mid-century periods (2041-2070) using the following equations (Campbell & Norman, 1998):

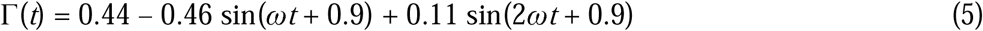

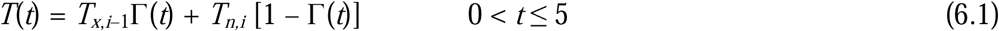

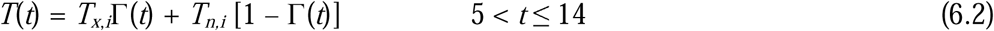

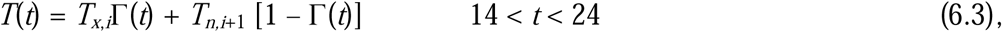

where ω = π/12; *t*: the time of day in hours (*t* = 12 at solar noon); *T_x_* and *T_n_*: daily maximum and minimum temperature, respectively, for the present day *_i_*, previous day *_i_*_–1_, and next day *_i_*_+1_.

Equation 5 is a dimensionless diurnal temperature function, while equations 6.1 – 6.3 comprise three similar formulas for the estimation of hourly temperature.

### 2.3. Calculation of chilling hours, chilling portions, and freezing events

The number of chilling hours and chilling portions for both the reference (1981-2000) and mid-century (2041-2070) periods were determined using hourly data. Daily and total accumulated chilling hours (Maulión et al., 2014)) and chilling portions (Fishman et al., 1987; Erez et al., 1989) were quantified. Chilling hours (32 °F < T < 45 °F) were calculated by counting the number of hours from 01 January to 31 December with estimated temperatures between 32 °F and 45 °F. For the determination of chilling portions (28.4 °F < T < 57.2 °F) (Erez et al., 1989), the estimated hourly temperature data was input into the Dynamic Model or Cumulative Chilling Portion model (https://ucanr.edu/site/fruit-nut-research-information-center/chilling-accumulation-modelsbr-their-calculation). Chilling portions are determined based on the Dynamic Model, which assumes that the degree of dormancy completion depends on the level of a certain dormancy-breaking factor, which accumulates in buds in a two-step process. The first step is assumed to be a reversible process that produces a thermally labile precursor. The formation of the precursor is promoted by chilling temperatures, while high temperatures reverse the process. Once the critical portion of the precursor is accumulated, it is transformed, irreversibly, in the second step to one portion of a stable dormancy-breaking factor or chilling portion (the unit of Dynamic Model). The derived constant values used in the program for the determination of chilling portions were: e0 = 4.15E+03; e1 = 1.29E+04; a0 = 1.40E+05; a1 = 2.57E+18; and slp = 1.6.

The daily probability of the occurrence of freezing events was also calculated using the cold sensitivity temperature for fully expanded corollas (T < −2.222 °C) (Smith ED, 2019). Historical freezing events were counted using T_min_ values from the Reference data set. The average number of days with T < −2.222 °C for each day (i.e., day 1-365) over the period assessed (20 years) was calculated from the local weather station data in Alma, GA. The projected likelihood of freezing events during the mid-century period (2041-2070) was determined by calculating T_min_ for each day (1/1/1981 – 31/12/2000) and GCM, using daily T_min_ (1/1/1981 – 31/12/2000) from the Reference data set and ΔTmin_m_ from each GCM. The number of predicted freezing events was averaged across all 20 GCMs for each day over the 20-year period to calculate the probability of freezing events for that day. For each of the same day (1-365), counts were averaged, resulting in the probability of freezing events for that day.

### 2.4. Growth chamber experimental design

The calculated temperature projections (i.e., hourly temperatures) for the reference (1981-2000) and mid-century (2041-2070) periods were used to design programs for a growth chamber experiment aimed at assessing the effects of ambient (i.e., historical) and elevated (i.e., projected) temperatures on blueberry. The set programs were operated in the ramping mode (a gradual and linear change in temperature over an hour period) (Table S1). Temperature and photoperiod were changed every 14 days according to the method described by Leisner et al. (2018). Data for photoperiod were obtained from the NOAA Solar Calculator (NOAA Global Monitoring Laboratory, 2022) using latitude 31.53580 and longitude −82.50670. Hourly temperatures and daily photoperiod were averaged in biweekly intervals starting on 01 October (Okie & Blackburn, 2011). Manual adjustments in the biweekly temperature dataset were necessary to compensate for the loss of chilling hours due to the averaging procedure. Light intensity was also controlled to mimic sunrise and sunset through a four-step increase or decrease, respectively, over a 1-hour period. Incandescent lamps were kept on during the entire daylength (Table S1). Temperature, relative humidity, and light intensity inside the growth chambers were recorded every 15 min using data loggers (Onset HOBO, Bourne, MA, USA). Relative humidity was set to ∼60% and the light intensity was set to reach 800 μmoles m^-2^s^-1^ when lights were on.

### 2.5. Plant material and temperature treatment

Three-year-old rabbiteye blueberry cultivars ‘Krewer’ and ‘Titan’ were obtained from Bottoms Nursery (Concord, GA, USA) on 01 December 2021. Plants were received in 1-gallon pots filled with pine bark and peat moss mixture. Plants were kept outside (Paterson Greenhouse Complex, Auburn, AL, USA Lat Lon) until 01 February 2022 when the first signs of breaking of dormancy appeared. When most of the plants had their floral buds at the bud swell stage, the plants were transferred into growth chambers (PGC-15, Percival Scientific, Perry, IA, USA) with controlled conditions as described above (Table S1). A total of ten plants (2 cultivars x 5 replicates) were placed in each of the growth chambers assigned for ambient and elevated temperature treatments. The temperature treatments in the growth chambers lasted until the late bloom stage (i.e., 8.5 weeks), with temperature and photoperiod from early February to early April simulated. The plants were watered throughout the duration of the experiment to maintain adequate soil moisture. Plants were rotated within the chamber every ∼2 weeks to ensure there were no within chamber effects.

### 2.6. Phenology observation and analysis

Data was collected for the Krewer and Titan cultivars for stages including the dates immediately after budbreak (BB), early bloom (EB), and late bloom (LB) observed inside the growth chambers under both ambient and elevated temperature conditions. For each plant in the ambient and elevated temperature treatments two to four buds from two to four branches were tagged upon transferring into the growth chambers. The floral buds on the tagged nodes were observed daily starting from initiation of the temperature treatment until all observed buds reached the late bloom stage. The stages were chosen due to their importance for management practices if they were in the field because they match the periods when major changes in phenology occur. Stages were defined based on the phenological stages chart for blueberries published by Michigan State University Extension (https://www.canr.msu.edu/blueberries/growing_blueberries/growth-stages). The dates of appearance for the key stages were analyzed using SAS (ver. 9.4; SAS Institute, Cary, NC) through descriptive statistics, Proc GLIMMIX, analysis of variance (ANOVA), and Fisher’s least significant difference (*p* < 0.05).

### 2.7. Model development

A nonlinear sigmoid model was adjusted to the observed values for the different stages over time using a logistic curve (Equation (7))

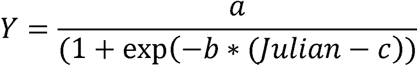

where *Y* is the percentage of each phenological stage, *a* is the maximum percentage of each stage *b* is the slope of the curve, and *c* is the value where the maximum rate of growth is reached in terms of Julian days. The daily rate of the percentage for each stage was derived from the logistic curve for each stage and ambient condition (average and elevated temperature).

## 3. Results and Discussion

### 3.1 Projected temperature changes for blueberry growing regions

Using the MACA data sets obtained for Bacon County, GA, the change in T_min_ and T_max_ (ΔTmin and ΔTmax) were determined by comparing the historical and projected temperatures during the reference (1981-2000) and mid-century (2041-2070) periods, respectively. A larger increase in maximum temperature compared to minimum temperature (i.e., ΔTmax > ΔTmin) was observed across all months except August and September for the mid-century (Supplemental Table S2; Table 1; Fig. 1). Both ΔTmin and ΔTmax were highest (2.9) during the month of August.

**Figure 1.**
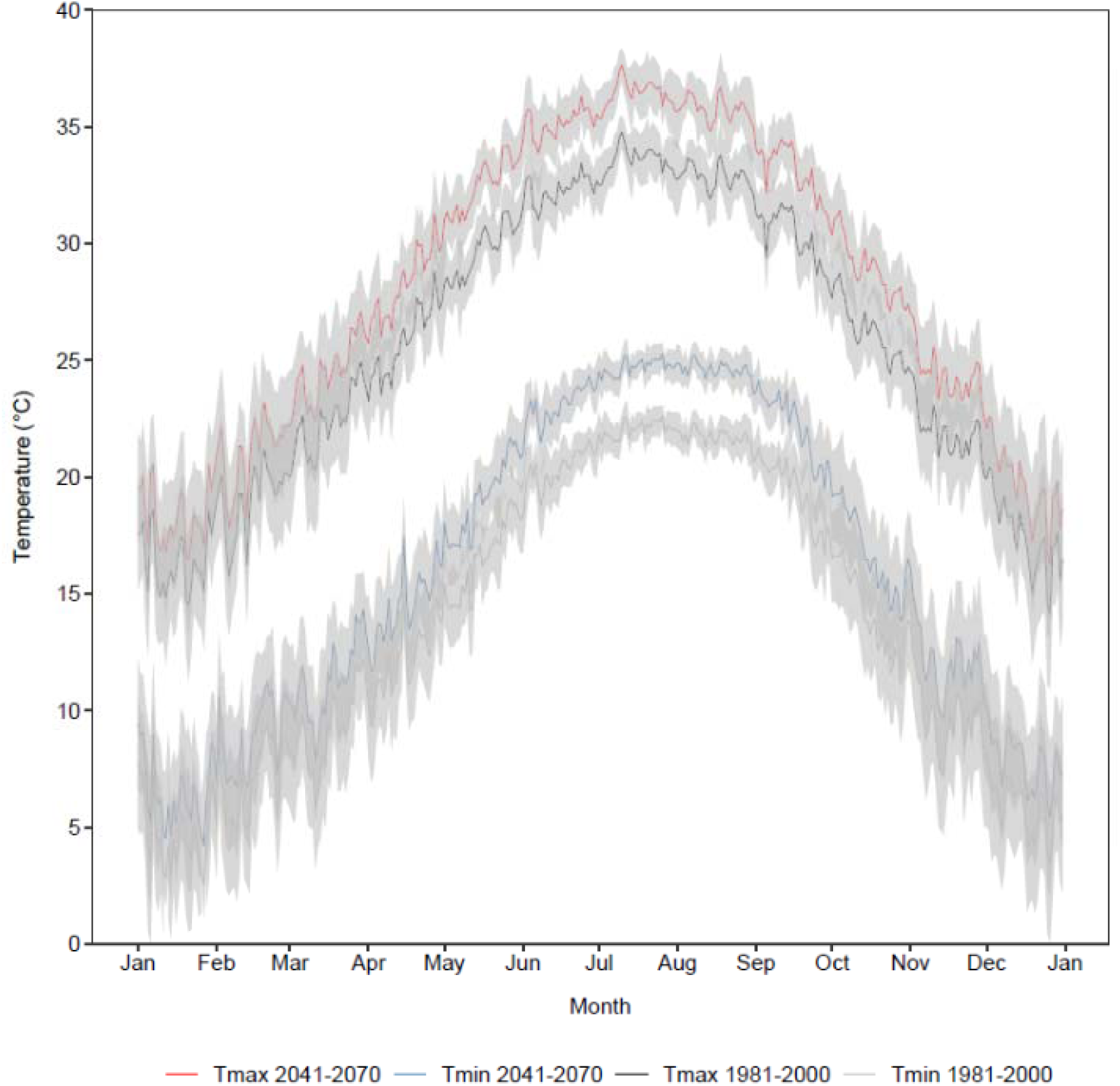
Daily minimum and maximum temperatures computed for the historical (1981 – 2000) and mid-century (2041 – 2070) period. The 95% confidence intervals (CI) are represented by the shaded area. For the estimation of CI for the historical period (1981-2000), daily minimum and maximum temperatures were calculated for each day (1/1 – 12/31) and year, and CI estimated across the 20 years. For the mid-century period (2041 – 2070), daily minimum and maximum temperatures were estimated for each day (1/1 – 12/31), year (1981-2000), and GCM. Daily minimum and maximum temperatures for each day were average across the 20 GCMs, and CI estimated across the 20 years.CI was estimated across the 20 years.

**Table 1.** Monthly ΔT_min_ and ΔT_max_ values. Data is computed on MACA data sets and represent the averages from 20 global climate models.

| Month | $\Delta T_{\min}$ | $\Delta T_{\max}$ |
| --- | --- | --- |
| January | 1.68 | 1.93 |
| February | 1.84 | 2.00 |
| March | 2.09 | 2.14 |
| April | 2.38 | 2.48 |
| May | 2.61 | 2.80 |
| June | 2.72 | 2.86 |
| July | 2.66 | 2.86 |
| August | 2.90 | 2.90 |
| September | 2.83 | 2.75 |
| October | 2.65 | 2.69 |
| November | 2.42 | 2.48 |
| December | 1.97 | 2.17 |

Conversely, the lowest ΔTmin (1.7) and ΔTmax (1.9) was recorded for January. The projected mean temperatures for the mid-century in this representative blueberry growing region ranged from 10.6 - 25.4 °C during the months of October to March (i.e., fall and winter seasons in the U.S.), which is the period when blueberry plants accumulate chill hours/portions. These projected temperatures are higher by 1.8-2.7 °C in comparison to the historical daily mean temperature (8.8-22.7 °C) for the October to March period.

### 3.2 Impact of projected elevated temperature on chill accumulation and freezing events

Using the projected temperatures for the mid-century period (2041-2070), the accumulation of chilling (32 °F < T < 45 °F) hours (Fig. 2; Supplemental Table S4) and chilling (28.4 °F < T < 57.2 °F) portions (Fig. 3) was quantified. Accumulation of chilling hours in the mid-century period is predicted to start on December 7^th^, four days later than in the reference period, and end on February 13^th^, 28 days earlier. The total amount of chilling hours to be accumulated in the mid-century period (167 hours) represents only 33.4% of the chilling hours accumulated in the historical period (500 hours). Moreover, the number of days in which accumulation of chilling hours occurs will decrease from 69 in the historical period to 32 in the mid-century. Higher differences in the number of days with accumulation between these periods will occur in December and February (Table S4).

**Figure 2.**
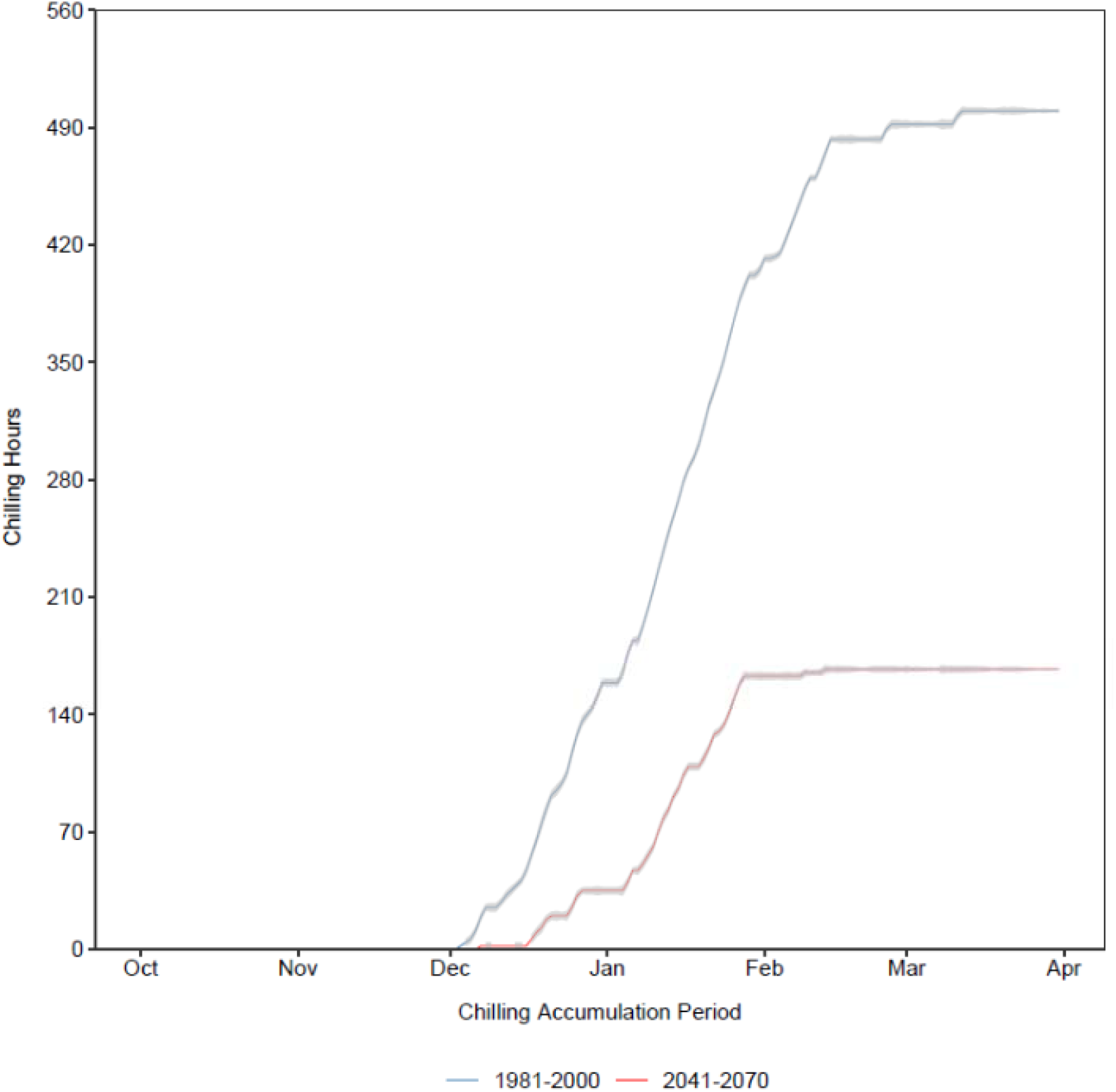
Accumulation of chilling hours (32 °F < T < 45 °F) for the historical (1981 – 2000) and mid-century (2041 – 2070) period. The 95% confidence intervals (CI) are represented by the shaded area. For the estimation of CI for the historical period (1981-2000), chilling hours were calculated for each day (1/1 – 12/31) and year, and CI estimated across the 20 years. For the mid-century period (2041 – 2070), chilling hours were estimated for each day (1/1 – 12/31), year (1981-2000), and GCM. The number of chilling hours for each day were average across the 20 GCMs, and CI estimated across the 20 years.

**Figure 3.**
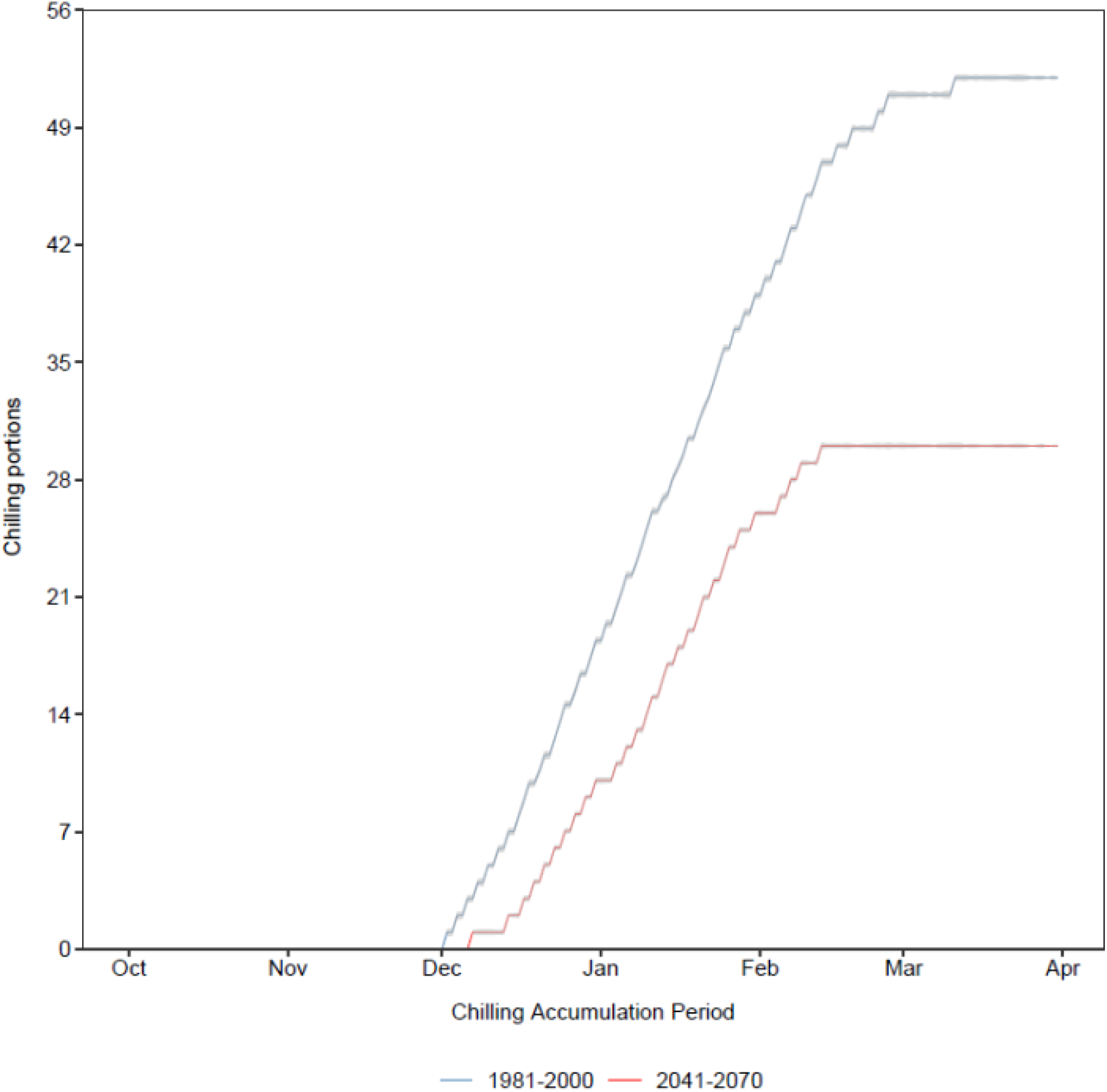
Accumulation of chilling portions (28.4 °F < T < 57.2 °F) for the historical (1981 – 2000) and mid-century (2041 – 2070) period. The 95% confidence intervals (CI) are represented by the shaded area. For the estimation of CI for the historical period (1981-2000), chilling portions were calculated for each day (1/1 – 12/31) and year, and CI estimated across the 20 years. For the mid-century period (2041 – 2070), chilling portions were estimated for each day (1/1 – 12/31), year (1981-2000), and GCM. The number of chilling portions for each day were average across the 20 GCMs, and CI estimated across the 20 years.

Chilling portions start to accumulate during the first week of December, but in the mid-century period it is predicted that accumulation of chilling portions will start five days later than the historical (1981-2000) period (Fig. 3). Moreover, it is projected that in the future, less chilling portions will be accumulated during October to March relative to historical data. The total chilling portions at the end of the accumulation period (31 March) in the middle of the century will be almost half (1.8x lower) than those accumulated during the historical period (Table S5). The projected mid-century temperatures also indicated that freezing events (T < - 2.222°C) are likely to occur within the chill accumulation period (October-March). While freezing events were first recorded around the middle of November during the historical period, this is expected to be delayed by eight days in the mid-century (Fig. 4). There were 84 days during the historical period that had a probability of freezing events that ranged from 5-30%. On the other hand, in the mid-century it is predicted that 77 days will have a 0.3-23.5% likelihood of experiencing freezing events. Consequently, days when freezing events occurred in the historical period (1981-2000) are also expected to experience this scenario in the future (2041-2070).

**Figure 4.**
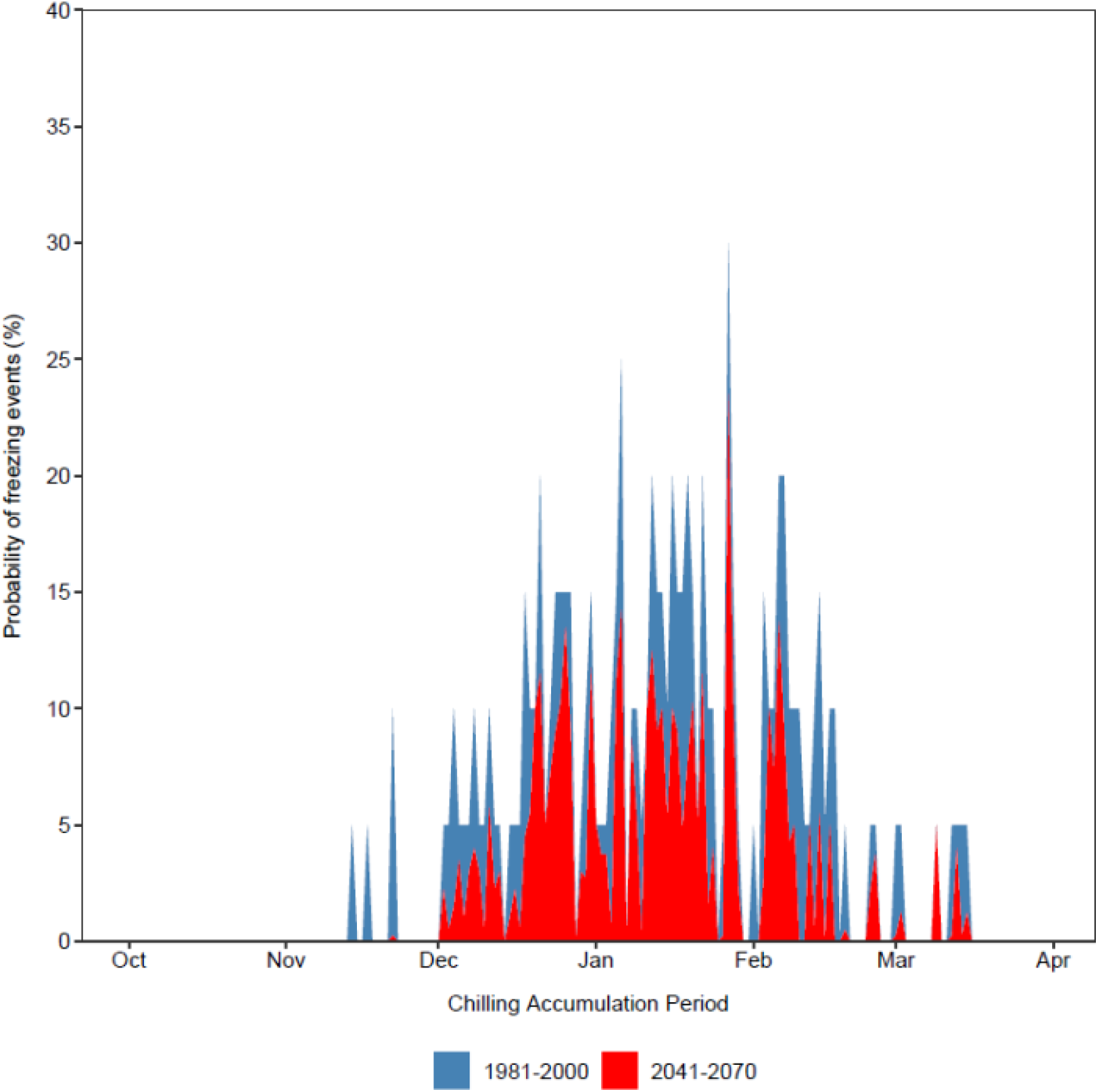
Probability of the occurrence of freezing events (T < −2.222 °C) for the historical (1981 – 2000) and mid-century (2041 – 2070) period.

Exception to this are two days in November (i.e., the time difference between the onset of freezing events) and five days in February. Aside from eight days that have the same probability of freeze events occurring in both periods assessed, all other days have a 1.0-12.3% lower probability in the mid-century relative to the reference years. Freezing events last until mid-March in both historical and mid-century periods.

Blueberry cultivars require varying levels of chilling for optimum flower and fruit development. Chilling accumulation hours (time spent at low temperatures) and forcing (accumulation of thermal time) are key traits associated with dormancy and dormancy release (Kovaleski, 2022). Out data suggested that plants need a longer period to accumulate chill hours/portions compared to historical data. This suggests that endodormancy will last longer and that dormancy release will occur later than usual. Lack of adequate chilling can negatively impact bud break and flowering time, which can lead to asynchronization of bloom time with pollinators, leading to floral bud death and yield impacts (Fadón & Rodrigo, 2018). Early release from dormancy can also make the plants more vulnerable to low temperatures in the spring (Salazar-Gutierrez et al., 2016). Our data indicates the likelihood of early spring frost events will increase in the mid-century period, having a potential negative impact on blueberry floral bud survival.

### 3.3 Effects of elevated temperature on blueberry phenology

Two cultivars of the rabbiteye ecotype (‘Krewer’ and ‘Titan’) were used to assess the effects of the future projected elevated temperature on the phenology of blueberry. The transition from dormancy to bloom was monitored. The onset of BB stage was significantly (*p* < 0.05 to *p* < 0.001) affected by all three factors analyzed (julian days, cultivar, and treatment (ambient and elevated temperature)), as well as by the interactions of cultivar with julian days and with treatment (Table 2). Similarly, for the EB stage, all single factors had a significant (*p* < 0.001) effect on the transition from late pink bud stage to early bloom (Table 3). Moreover, except for the julian days x treatment interaction, all two- and three-factor interactions also significantly (*p* < 0.05 to *p* < 0.001) affected the onset of the EB stage. Lastly, all three factors and their interactions significantly (*p* < 0.05 to *p* < 0.001) affected the transition from the EB to LB stage (Table 4).

**Table 2.**
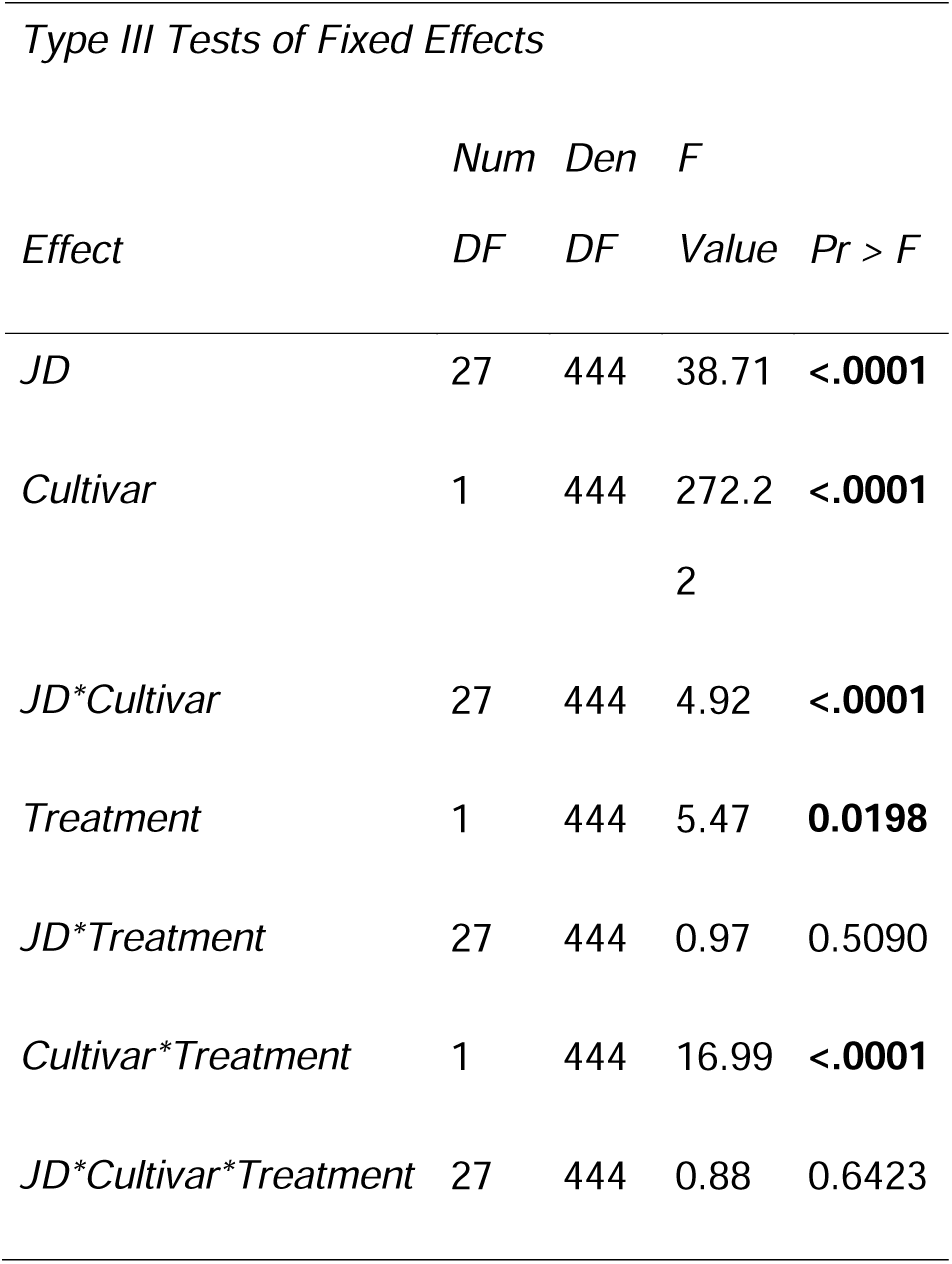
ANOVA results for the effect of projected elevated temperature on BB stage. JD = julian day. *P* values in bold imply significant effect of the factor.

**Table 3.**
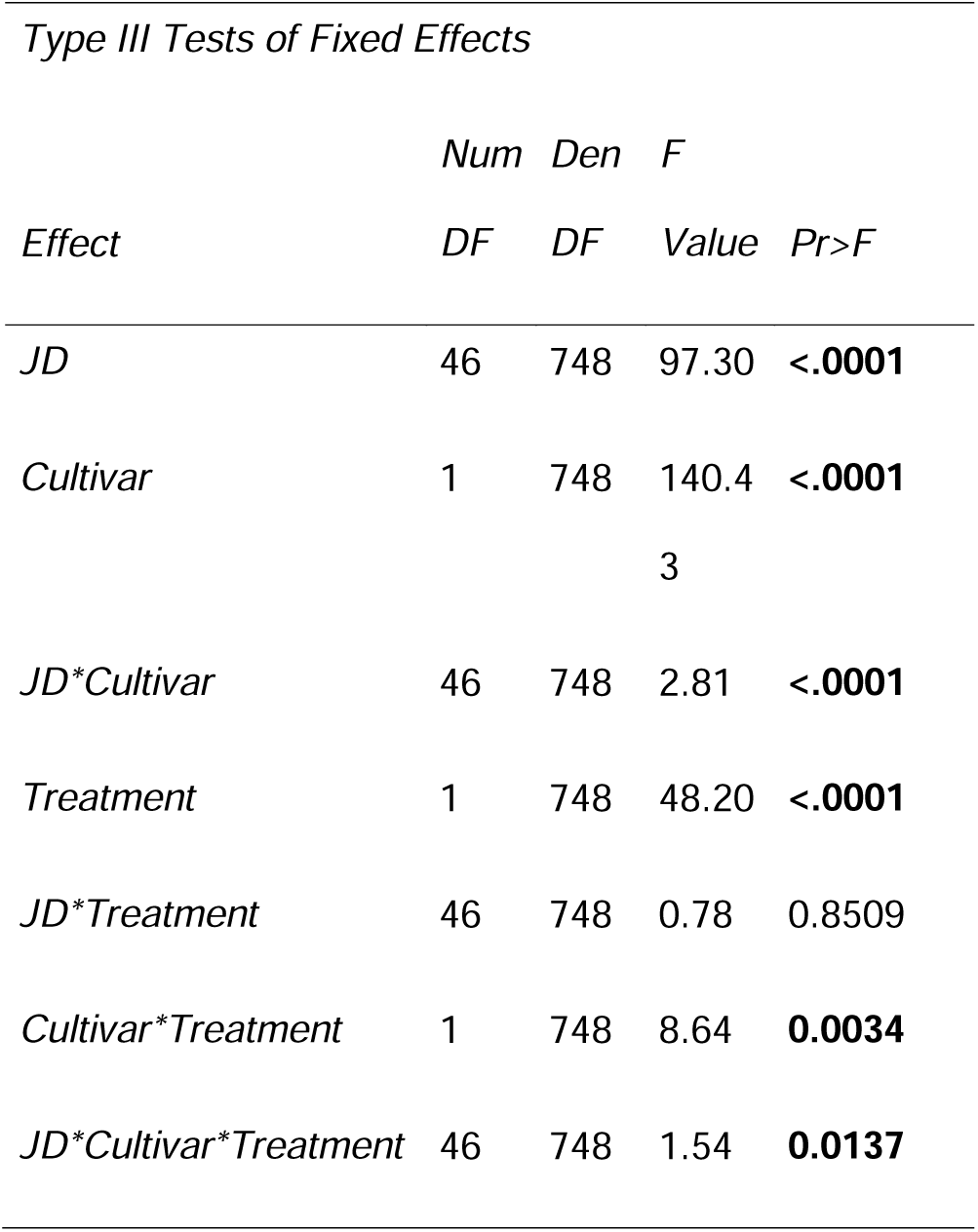
ANOVA results for the effect of projected elevated temperature on EB stage. JD = julian day. *P* values in bold imply significant effect of the factor.

**Table 4.**
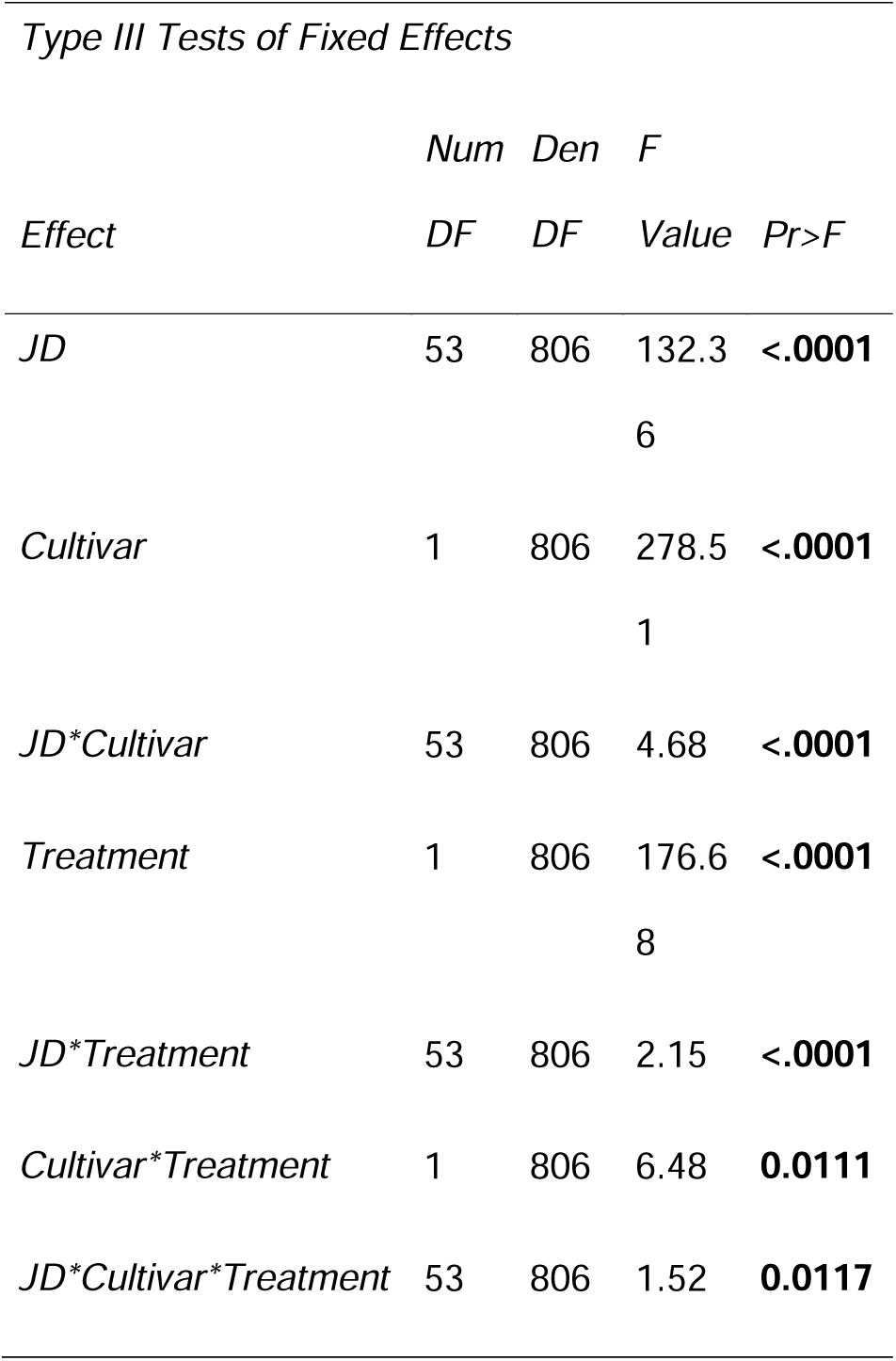
ANOVA results for the effect of projected elevated temperature on FB stage. JD = julian day. *P* values in bold imply significant effect of the factor.

The impact of the temperature on three key phenological stages including BB, EB, and LB, was modeled and the rate in terms of percentage for each stage was estimated (Figure 5). Parameters were estimated for each cultivar (Table 5), condition and phenological stage. It was observed that for BB stage even though all the cultivars reached the maximum percentage the duration was different between ‘Krewer’ (K) and ‘Titan’(T) cultivars under ambient (A) vs. elevated (E) conditions. Titan under elevated conditions (TE) (Figure 5A,B) presented the steepest curve of the accumulated percentage and the maximum daily rate was reached at 50 (DOY) in contrast with ‘Titan’ under ambient conditions (TA) where the lowest rate was observed and the accumulated percentage took more days to reach the 100%. ‘Krewer’ started early on both conditions (KA, KE) reaching the maximum percentage almost at the same time as TA. For all stages the duration to reach the maximum percentage of any stage ranges between 41 to 94 days. For the EB stage (Figure 5 C,D) the only cultivar that did not reached 100% of the first bloom was KE. TA started first followed by KE, KA, and TE that for this stage took more days to reach the maximum percentage. However, TE presented the highest daily rate to reach the stage followed by KA KE and TA. For the last stage LB, (Figure 5 E,F), TA did not reach 100%. KE started followed by KA and TE that crossed around the day 75 and the last to reach the stage was TA. In terms of daily rate TE presented the highest daily rate followed by KE, KA, and TA. Overall, TE was constant in presenting the highest daily rate for all stages while the lowest rate was observed on TA.

**Figure 5.**
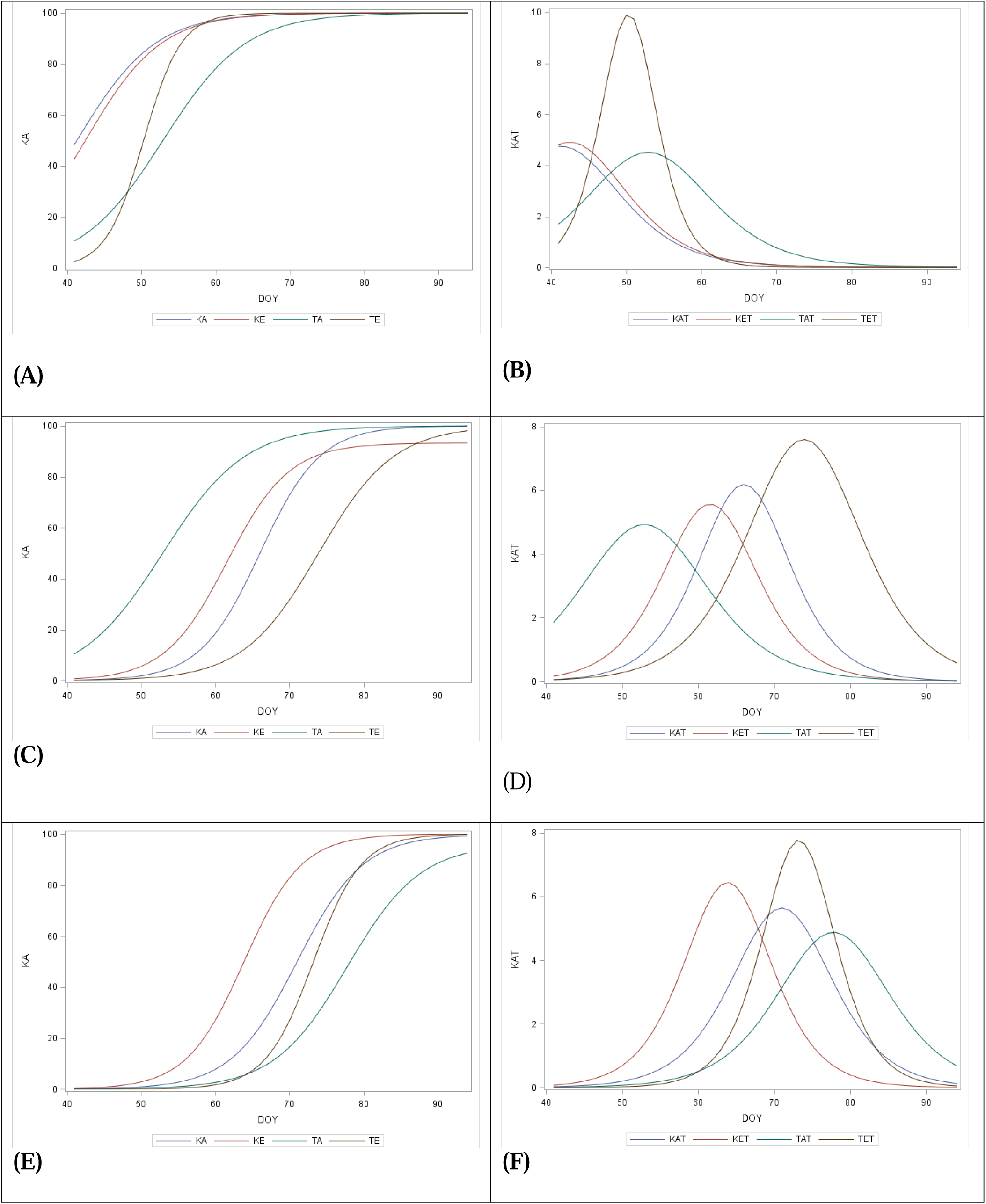
Models for ‘Krewer’(K) and ‘Titan’ (T) (left column) and rates (right column) for the duration (DOY) and percentage (KA and KAT) to reach bud break (A,B), early bloom (C,D), and late bloom (E,F) under ambient (A) and elevated temperatures (E).

**Table 5.**
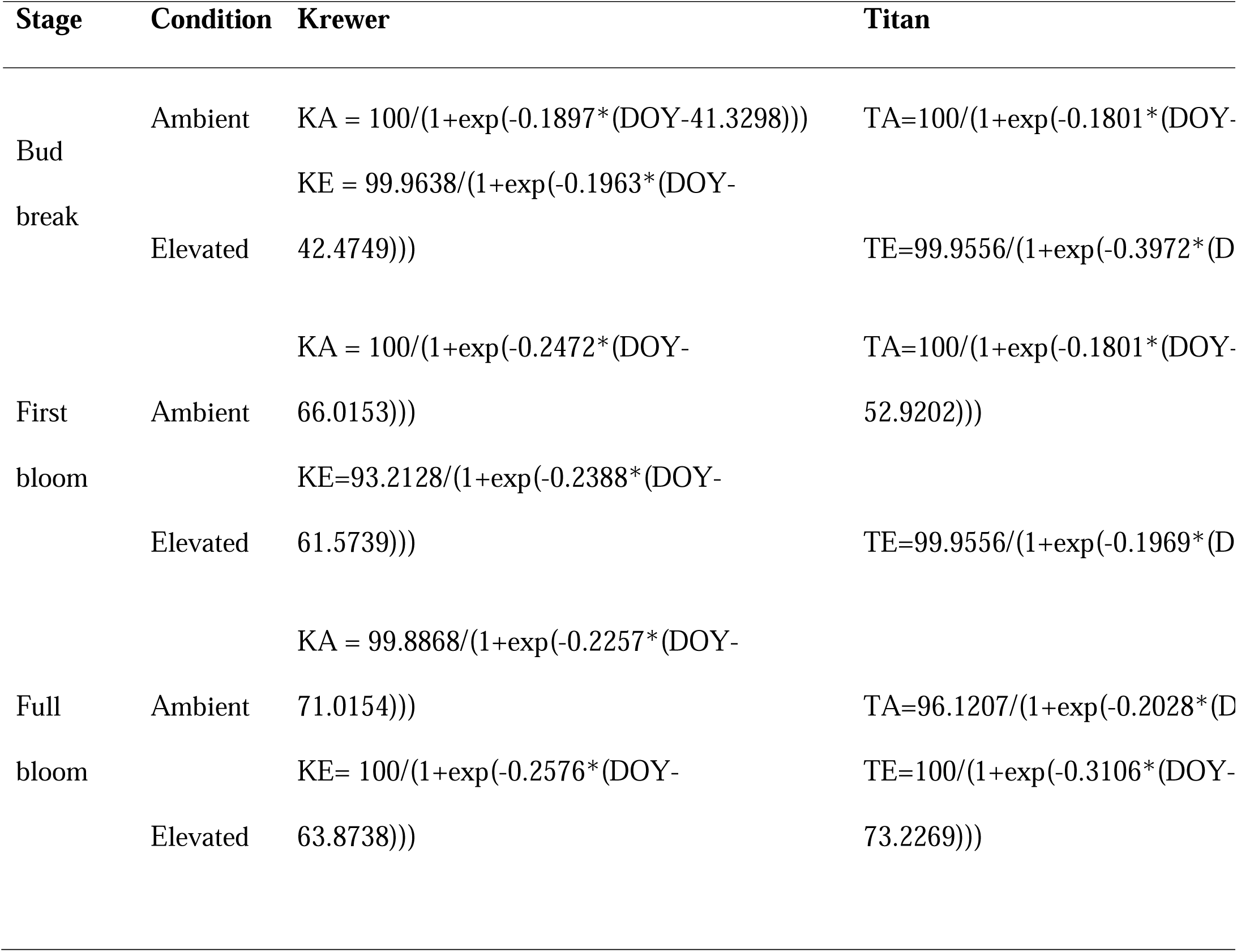
Models adjusted for three key phenological stages for cultivars ‘Krewer’(K) and ‘Titan’(T) for each condition ambient (A) and elevated (E).

## 4. Conclusions

Our analysis showed that by the middle of the century, temperatures will be higher relative to the historical/reference period of 1981-2000. This warming is predicted to reduce the chill hours/portions that plants can accumulate, which could negatively affect their development.

Despite the increased temperatures in the future, freezing events are still expected to occur, which could negatively affect plant productivity depending on the stage which it coincides. Furthermore, we generated a model explaining the impacts of elevated temperature on blueberry phenology, in particular bud break and bloom. This model would be useful in predicting bud break and bloom dates under future climates. These findings underscore the physiological sensitivity of blueberry dormancy to climate change and highlight the need for a deeper understanding of the genetic and environmental factors involved.

## Acknowledgments

We thank Dr. Jay Spiers for his help in acquiring the blueberry plants and recommendations for crop husbandry. This work was supported by a grant from the Alabama Agricultural Extension System (1018601) to CPL and the USDA NIFA Award (No. 2022-67013-41879) to CPL.

